# Phylogeography of the Giant Honeybees Based on Mitochondrial Gene Sequences

**DOI:** 10.1101/2024.03.30.587434

**Authors:** Chet P. Bhatta, Sarah C. Cluff, Deborah R. Smith

## Abstract

We carried out a phylogenetic analysis of the giant honeybees using mitochondrial COI and COII gene sequences analyzed with maximum likelihood methods. Our goal was to resolve phylogenetic relationships among *Apis laboriosa*, and the *Apis dorsata* subspecies: *A. d. dorsata, A. d. binghami*, and *A. d. breviligula*, the last two of which have been proposed as full species by several authors. We obtained strong support for four clades within *A. dorsata*: the three subspecies mentioned above, and a fourth from south India, but our analysis did not resolve the phylogenetic relationships among the four clades within *A. dorsata* in the broad sense. Recognition of these distinct lineages is important for conservation planning, so that their individual ecologies and migration patterns can be taken into account.

## Introduction

The giant honeybees have a range centered on south and southeast Asia, extending northwest into Pakistan, eastwards through India, Bangladesh, Nepal, Bhutan, Myanmar, southern China, and southeast Asia, and through the islands of Malaysia, Indonesia, and the Philippines. Several earlier writers including Maa (1953) and Ruttner (1988) pointed out geographical diversity among giant honeybee populations. In particular they noted that the giant bees of the Himalayan region, the Indonesian island of Sulawesi, and the oceanic Philippine islands (i.e., those never connected to the Asian mainland) differed from one another and from the more widespread form found elsewhere. Maa divided honeybees into three genera—*Micrapis*, the dwarf honeybees, *Megapis*, the giant honeybees, and *Apis*, the cavity nesting honeybees—and recognized four giant bee species: *Megapis breviligula* from the Philippines, *M. binghami* from Sulawesi and smaller nearby islands, *M. laboriosa* from high altitude Himalayan regions, and the more widespread *M. dorsata*. Ruttner, like most subsequent authors, recognized just one genus, *Apis*, and only one species of giant honeybee, *Apis dorsata*. He and many subsequent authors (e.g., Engel, 1999, 2002) considered the Himalayan form a subspecies, *A. d. laboriosa*, but noted that additional information might confirm it as a distinct species.

The species status of *A. laboriosa* remained contentious for many years despite numerous studies. Sakagami et al. (1980) made detailed morphological comparisons of *A. laboriosa* from Nepal and *A. dorsata* collected from many parts of its range, documenting “distinct and stable differences between them” supporting species status of *A. laboriosa*. McEvoy and Underwood (1988) reported that they could find no morphological differences between male genitalia (the everted endophallus) of *A. laboriosa* and *A. dorsata*, but nonetheless supported species status of *A. laboriosa* on the basis of other morphological differences, habitat, the presence of two species of braulid parasites (Diptera: Bruaulidae, *Megabraula*) in nests of *A. laboriosa* but (apparently) not those of *A. dorsata*, and genetic differences revealed by allozyme electrophoresis.

However, some authors argued that the characters used to support species status of *A. laboriosa*—including habitat, color patterns, and morphometric characters—could represent intraspecific variation and adaptation to different habitats, and thus took the conservative position that more data were needed, particularly concerning reproductive isolation of populations occurring in sympatry (e.g., Ruttner, 1988, Engel, 1999). Cao et al. (2012a) carried out morphometric comparisons of *A. laboriosa* and *A. dorsata* collected from Yunnan, Guangxi and Hainan provinces in China and again found significant differences between them. Collection sites for the two were in relatively close proximity (on the order of 200-300 km) but not strictly sympatric, and they were found at different elevations (*A. laboriosa* 1500 m and above, *A. dorsata* 1300 m and lower, though all but one collection was made at 700 m or lower).

More recently, new distributional records for *A. laboriosa* (Kitnya et al., 2020) reported *A. dorsata* and *A. laboriosa* foraging sympatrically at sites in Arunachal Pradesh, India and in northern Vietnam. Kitnya et al. (2022), found distinct morphological, morphometric, and genetic differences between Indian populations of *A. dorsata* and *A. laboriosa*, both in sympatry and in allopatry, providing convincing support for the species status of *A. laboriosa*.

Until recently the species status of *A. d. breviligula* and especially of *A. d. binghami* have received much less attention. Arias and Sheppard (2005) included A. *laboriosa, A. dorsata* from Thailand and Sri Lanka, and *A. binghami* in a larger phylogenetic analysis of *Apis* species using both nuclear (EF-1α intron) and mitochondrial (ND2) sequence data. The giant bees were recovered as a monophyletic group and *A. laboriosa* was consistently recovered as a clade distinct from *A. dorsata* and *A. d. binghami*; however, the latter two were not consistently resolved as separate lineages. Raffiudin and Crozier (2007) used both mitochondrial (cytochrome c oxidase 2, ND2, and the large (16S) ribosomal subunit or *rrnL*) and nuclear (inositol 1,4,5-triphosphate receptor or *itpr*) gene sequences in their phylogenetic analysis of *Apis* taxa, also including the giant bees *A. dorsata, A. d. binghami*, and *A. laboriosa*. Their analyses consistently recovered *A. laboriosa* as sister to *A. dorsata* and *A. d. binghami*. Lo et al. (2009) carried out a more comprehensive coverage of giant bees, including *A. laboriosa, A. dorsata, A. d. binghami* and *A. d. breviligula* in their phylogenentic analysis of *Apis* species, also the same sets of gene as Raffiudin and Crozer (2007) minus the mitochondrial ND2. Their results strongly supported the species status of *A. d. breviligula* from the Philippines, though the placement of *A. d. binghami* remained unresolved.

In this paper, we accept the species status of *A. laboriosa*. We use the names *A. dorsata dorsata, A. d. breviligula* and *A. d. binghami* for the other distinctive populations of giant honeybees because the species status of the latter two taxa, particularly A. d. binghami, is still subject to investigation. We use the name “A. d. South India” to refer to a population that appears to be a cryptic unnamed species or subspecies (Smith, 1991, Kitnya et al., 2022). “*Apis dorsata* in the broad sense” will refer to all giant bees excluding *A. laboriosa*.

The objective of this study is to construct a phylogenetic tree for populations of giant honeybees, including representatives from as much of their range as we could obtain, to test whether the lineages within *A. dorsata* in the broad sense are monophyletic, and to determine relationships among them. Samples include *A. laboriosa*, multiple populations of *A. d. dorsata*, and the distinctive island populations of Sulawesi and smaller nearby islands (*A. d. binghami*) and the oceanic Philippines (*A. d. breviligula*). We also include the dwarf bees *A. florea* and *A. andreniformis* and the cavity-nesting bees *A. mellifera* and *A. cerana* as outgroups. We generated partial sequences of the mitochondrial cytochrome c oxidase subunit 1 (COI) and cytochrome c oxidase subunit 2 (COII) genes and used Maximum Likelihood methods in MEGA7 to construct phylogenetic trees.

## Methods

### Field Methods

Samples used in this study were collected by multiple researchers from 1989 to 2018 using a variety of collection and preservation techniques. Table 1 gives locality and collection information, and sample IDs corresponding to those used in Figure 1. Most specimens were collected directly from colonies, though some bees were collected while they were foraging. Most specimens are adult worker bees, while a few are pupae collected directly from nests. Individual bees or bee thoraces were preserved in the field in liquid nitrogen (1988-1990) or in 95% ethanol (1991 onwards). Frozen specimens were later stored at -80°C. Ethanol-preserved specimens were stored at 4° to -20C.

**Table 1.** Bee samples used in this study. Taxa and DNA ID code are used in the phylogenetic tree presented in the Figure 1. More detailed collection information is presented in Supplemental File 1.

| Taxa | ID code (see Figure 1) | Country | Locality | Genbank Accession # |
| --- | --- | --- | --- | --- |
| ***Apis laboriosa*** |  |  |  |  |
| *A. laboriosa* | DNA-13232 NEPAL | Nepal | Baglung | xxxx |
| *A. laboriosa* | DNA-13233 NEPAL | Nepal | Kaski | xxxx |
| *A. laboriosa* | DNA-13235 NEPAL | Nepal | Kaski | xxxx |
| *A. laboriosa* | GB-AP018039 NEPAL | Nepal | unknown | AP018039 |
| ***Apis dorsata dorsata*** |  |  |  |  |
| *A. d. dorsata* | DNA-14b PAKISTAN | Pakistan | Islamabad | xxxx |
| *A. d. dorsata* | DNA-12956-57 NEPAL | Nepal | Kanchanpur | xxxx |
| *A. d. dorsata* | DNA-12958-59 NEPAL | Nepal | Kanchanpur | xxxx |
| *A. d. dorsata* | DNA-13129 NEPAL | Nepal | Kohalpur | xxxx |
| *A. d. dorsata* | DNA-13131 NEPAL | Nepal | Bardiya | xxxx |
| *A. d. dorsata* | DNA-BPO18 INDIA- Assam | India (northeast) | Assam | xxxx |
| *A. d. dorsata* | DNA-BPO19 INDIA- Assam | India (northeast) | Assam | xxxx |
| *A. d. dorsata* | DNA-1a INDIA-Andaman Is. | India | Andaman Is. | xxxx |
| *A. d. dorsata* | DNA-HP88-N09THAILAND-Chiangmai | Thailand | Chiang Mai x | xxxx |
| *A. d. dorsata* | GB-AP018369 THAILAND-Bangkok | Thailand | Bangkok | AP018369 |
| *A. d. dorsata* | DNA-GWO89-07 MALAYSIA-peninsula | Malaysia | Peninsula | xxxx |
| *A. d. dorsata* | DNA GWO89-97A MALAYSIA-Borneo | Malaysia | Borneo | xxxx |
| *A. d. dorsata* | DNA-12906 INDONESIA-Timor | Indonesia | Timor | xxxx |
| *A. d. dorsata* | DNA-12907 INDONESIA-Timor | Indonesia | Timor | xxxx |
| *A. d. dorsata* | DNA-12910 INDONESIA- Flores | Indonesia | Flores | xxxx |
| *A. d. dorsata* | DNA-SR92-007 PHILIPPINES- Palawan | Philippines | Palawan Island |  |
| ***Apis dorsata (South India)*** |  |  |  |  |
| *A. dorsata*-SIndia | SI DNA-08b S-INDIA | India (south) | Karnataka | xxxx |
| *A. dorsata*-SIndia | SI DNA-12127 S-INDIA | India (south) | Tamil Nadu | xxxx |
| *A. dorsata*-SIndia | SI DNA-12128 S-INDIA | India (south) | Karnataka | xxxx |
| *A. dorsata*-SIndia | SI DNA-12129 S-INDIA | India (south) | Tamil Nadu | xxxx |
| *A. dorsata*-SIndia | SI DNA-10240 S-INDIA | India (south) | Karnataka | xxxx |
| ***Apis dorsata breviligula*** |  |  |  |  |
| *A. d. breviligula* | DNA-12905 PHILIPPINES- Mindanao | Philippines | Mindanao Island | xxxx |
| *A. d. breviligula* | DNA-13217 PHILIPPINES- Luzon | Philippines | Luzon Island | xxxx |
| *A. d. breviligula* | DNA-13222 PHILIPPINES- Luzon | Philippines | Luzon Island | xxxx |
| ***Apis dorsata binghami*** |  |  |  |  |
| *A. d. binghami* | DNA-12912 INDONESIA-Sulawesi | Indonesia | S. Sulawesi | xxxx |
| *A. d. binghami* | DNA-GWO89-81B INDONESIA-Sulawesi | Indonesia | S. Sulawesi | xxxx |
| *A. d. binghami* | DNA-GWO89-80A INDONESIA-Sulawesi | Indonesia | S. Sulawesi | xxxx |
| **Short Sequences not included in phylogenetic analysis** |  |  |  |  |
| *A. d. dorsata* |  | India | Mizoram | KU212344.1 |
| *A. d. dorsata* |  | India | Mizoram | KU212345.1 |
| *A. d. dorsata* |  | Myanmar |  | MFBO4562.1 |
| *A. d. dorsata* |  | Myanmar |  | MF804563.1 |
| **Outgroups** |  |  |  |  |
| *A. mellifera ligustica* | ligustica L06178.1 | Australia |  | L06178.1 |
| *A. cerana* | NC014295 CHINA | China |  | NC014295 |
| *A. andreniformis* | DNA-4668 THAILAND | Thailand | Surat Thani | xxxx |
| *A. andreniformis* | DNA GWO89-113 MALAYSIA-Borneo | Malaysia | Borneo | xxxx |
| *A. florea*- East | DNA-10435 THAILAND | Thailand | Ratchabuti | xxxx |
| *A. florea*- East | DNA-10434 THAILAND | Thailand | Ratchaburi | xxxx |
| *A. florea*- East | DNA-7014 CAMBODIA | Cambodia | Kampong Spoe | xxxx |
| *A. florea*- East | DNA-10246 CAMBODIA | Cambodia | Kampong Spoe | xxxx |
| *A. florea*- East | DNA-10248 CAMBODIA | Cambodia | Kampong Spoe | xxxx |
| *A. florea*- West | DNA-9744 SAUDI ARABIA | Saudi Arabia | Hasa | xxxx |
| *A. florea*- West | DNA-COLONY 1 ISRAEL | Israel | Eilat | xxxx |
| *A. florea*- West | DNA-10264 INDIA | India (south) | Karnataka | xxxx |
| *A. florea*- West | DNA-10265 INDIA | India (south) | Karnataka | xxxx |
| *A. florea*- West | DNA-10222 INDIA | India (south) | Karnataka | xxxx |
| *A. florea*- West | DNA-10224 INDIA | India (south) | Karnataka | xxxx |

**Figure 1.**
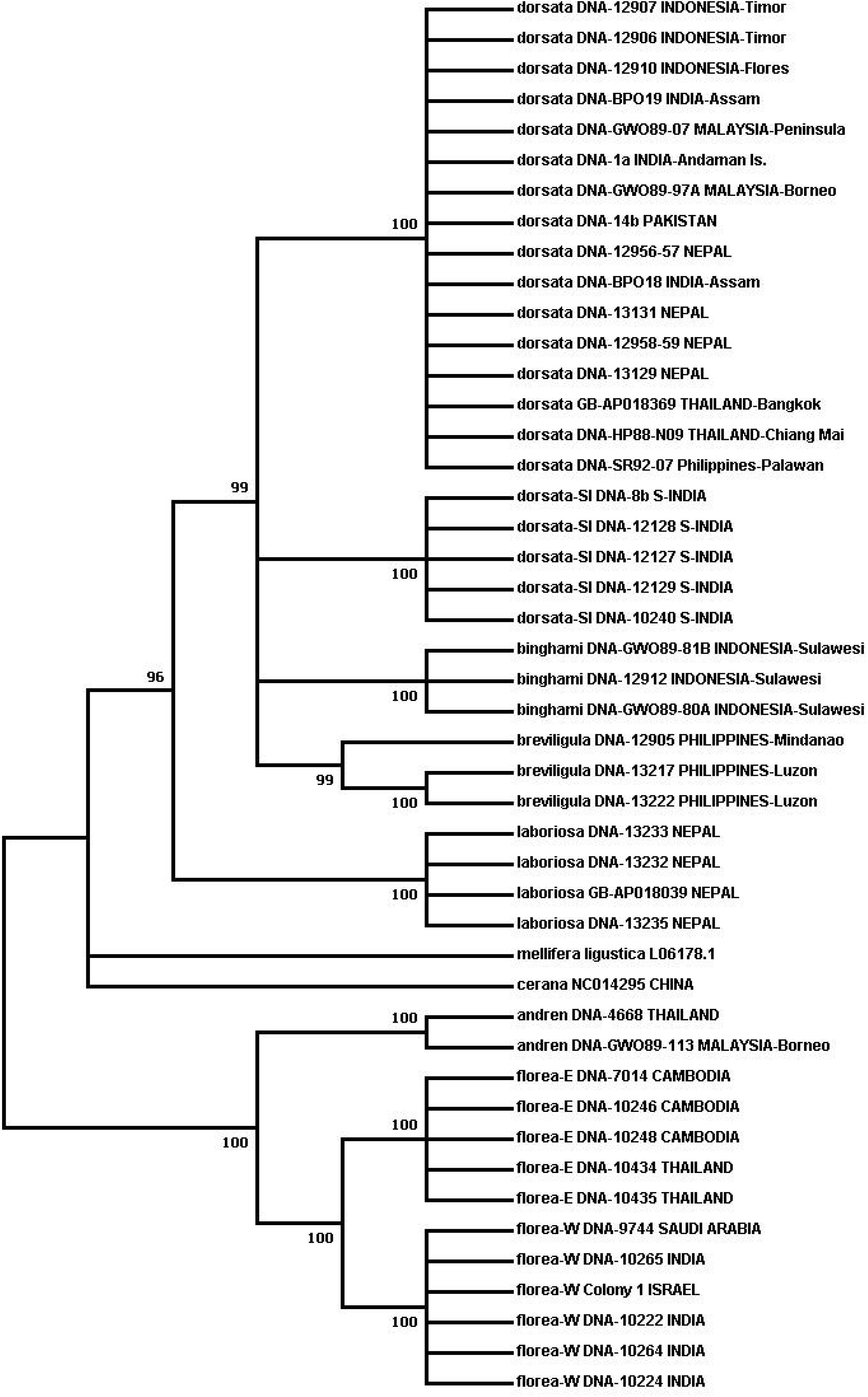
Phylogenetic tree obtained using the Maximum Likelihood method based on the General Time Reversible model. Numbers on branches indicate bootstrap support; partitions with less than 95% support were collapsed. See text for more detailed information on analysis methods.

### Laboratory methods

Genomic DNA was extracted from the mitochondrion-rich thoracic flight muscle tissue using DNA spin-columns, primarily the Qiagen DNEasy Blood and Tissue kit (www.Qiagen.com) and the GenElute Mammalian Genomic DNA Miniprep kit (www.sigmaaldrich.com) following the manufacturers’ recommendations. Extracted DNAs were stored at -20°C. Portions of the mitochondrial genome were amplified using the primers shown in Table 2. These sequences included a large portion of COI, leucine tRNA_UUR_, a short non-coding sequence, and a portion of COII. Figure 2 shows the relative position of the primers on the honeybee COI-COII sequences. Sanger sequencing was carried out at the Idaho State University Molecular Research Core Facility, Pocatello, ID. As only protein-coding sequence was included in the phylogenetic analysis, the tRNA and non-coding sequences were removed after alignment (see below) and the COI and COII sequences were concatenated. Some of the sequences were also obtained from Genbank (Table 1). The total number of sequences for each taxon and their geographic origins is summarized in Table 3.

**Table 2.**
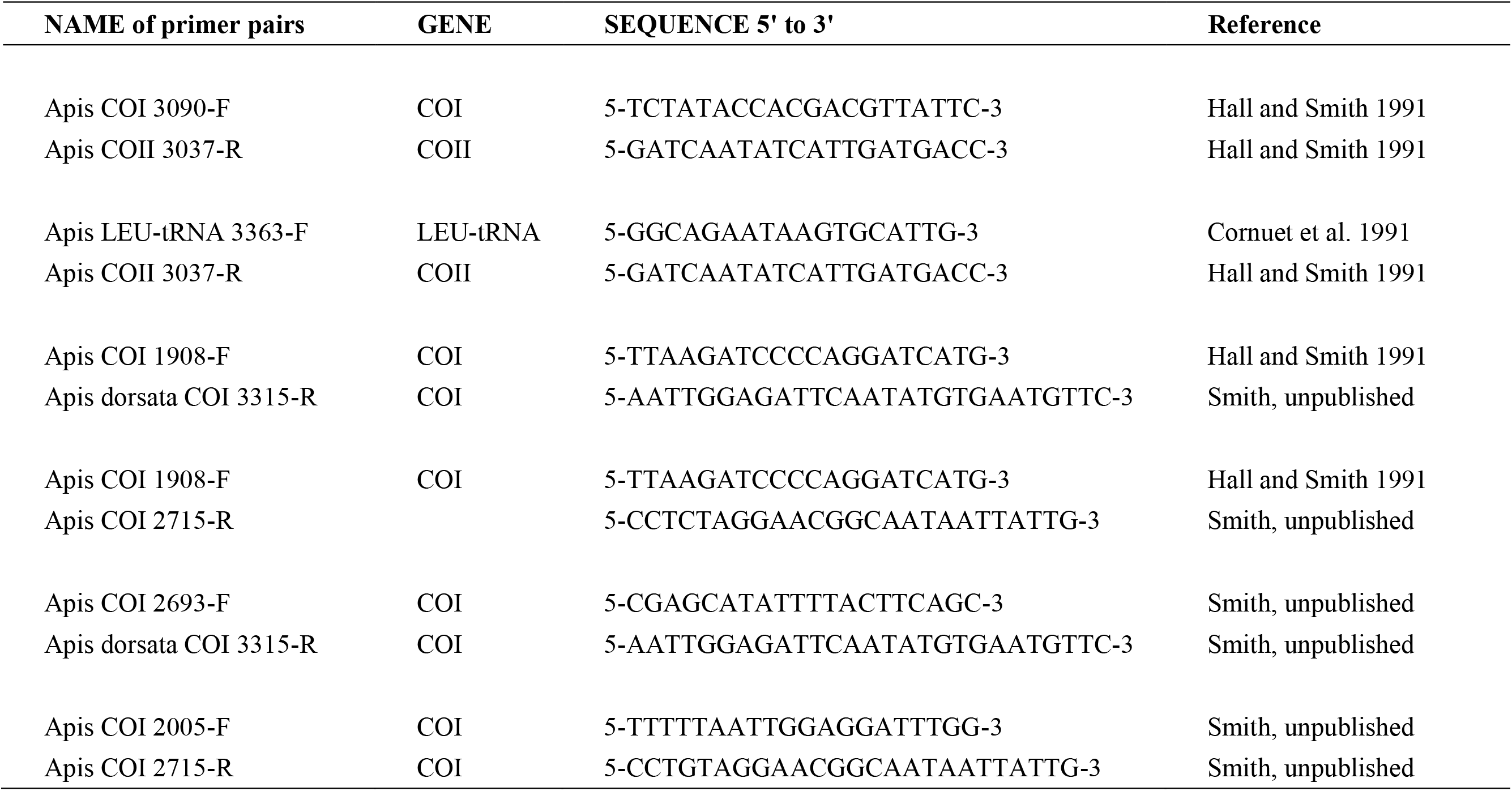
PCR primer sequences used in this study. “GENE” indicates the gene the primer bids to. Numbers in the primer name refer to the position of the 5’ end of the primer on the complete mitochondrial genome of *Apis mellifera ligustica* (Genbank Accession #L06178.1; Crozier and Crozier 1993).

**Table 3.** Summary table showing number of sequences for each taxon and their geographic origins.

| Species | # sequences | Country of Origin |
| --- | --- | --- |
| **Giant bees** |  |  |
| *A. laboriosa* | 4 | Nepal |
| *A. d. dorsata* | 16 | Pakistan, India (northeast, Andamans), Nepal, Thailand, Malaysia (peninsula and Borneo), Indonesia, Philippines (Palwan) |
| *A. d. South India* | 5 | India (south) |
| *A. d. breviligula* | 3 | Philippines |
| *A. d. binghami* | 3 | Indonesia (Sulawesi) |
| **Outgroups** |  |  |
| *A. andreniformis* | 2 | Thailand, Malaysia (Borneo) |
| *A. florea- East* | 5 | Cambodia, Thailand |
| *A. florea- West* | 6 | India, Saudi Arabia, Israel |
| *A. mellifera ligustica* | 1 | Australia |
| *A. cerana* | 1 | China |

**Figure 2.**
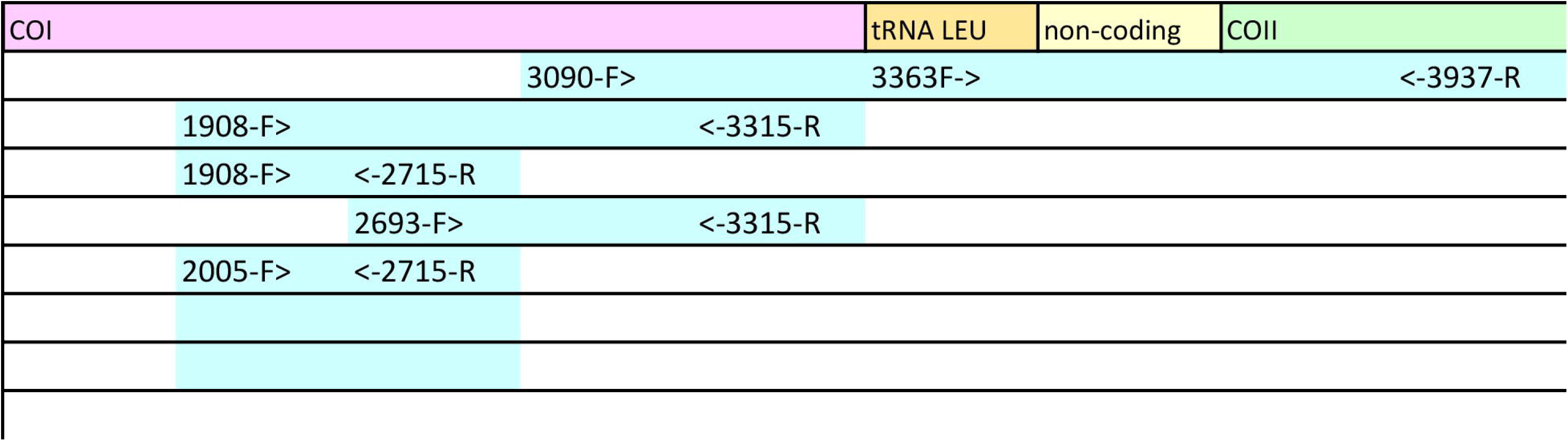
Relative position of primers on the mitochondrial COI, COII and leucine tRNA genes. Numbers in the primer name refer to the position of the 5’ end of the primer on the complete mitochondrial genome of *Apis mellifera ligustica* (Genbank Accession #L06178.1; Crozier and Crozier 1993). Not drawn to scale.

### Phylogenetic analysis

Sequences were aligned manually with COI and COII sequences from *Apis mellifera ligustica* (Crozier and Crozier,1993; Genbank accession L06178) in MEGA7 (Kumar et al., 2016). Sequences were screened for missing bases and correct reading frame by translating DNA sequences to amino acid sequences. In total, 46 sequences were used in the phylogenetic analysis and another four Genbank sequences of *A. dorsata dorsata* from India (Mizoram) and Myanmar (Table 1) that were too short to include in the phylogenetic analysis were aligned with the larger data set to determine which sequences they matched most closely.

The best model of sequence evolution was selected using MEGA “Model Selection” analysis and the following conditions: maximum likelihood statistical methods, partial deletion of sites with missing data, coverage cutoff of 75%, all codon positions used, moderate branch swapping filter. The model of sequence evolution with the lowest Bayesian Information Criteria (BIC) score was selected for use in the phylogenetic analysis. This model (BIC score 13302.38) was a general time reversible model with non-uniform rates of evolution among sites (gamma distributed) and a fraction of sites seemingly invariable (GTR+G+I).

Phylogenetic trees were constructed using Maximum Likelihood methods in MEGA7 with the following settings: model of evolution gamma distributed with invariant sites (GTR+G+I) with 5 gamma categories, partial deletion of sites with missing data, 75% site coverage cutoff, all codon positions used, maximum likelihood heuristic method Subtree-Pruning-Regrafting - Fast, initial tree generated by Neighbor Joining, moderate branch swap filter, 3 threads. Support for the branching patterns was evaluated with 1000 bootstrap replicates.

Branches with less than 95% bootstrap support were collapsed. A coverage cutoff of 75% was chosen during model choice and tree-building to ensure that inclusion of shorter sequences did not result in elimination of informative data.

## Results

Figure 1 presents the phylogenetic tree obtained showing partitions with 95% bootstrap support or better. As has been found in other recent studies, *A. laboriosa* constitutes a well-supported lineage (100% bootstrap support) separate from and sister to all *A. dorsata* in the broad sense, further supporting its status as a distinct species. Within *A. dorsata* in the broad sense, we found four distinct lineages: *A. d. breviligula* from the oceanic Philippine islands (99% bootstrap support), *A. d. binghami* from the Indonesian island of Sulawesi (100% bootstrap support), A. d. South India, a genetically distinct southern Indian population (100% bootstrap support), and a more narrowly defined *A. dorsata dorsata*, represented by our samples from Pakistan, Nepal, northeastern India (Assam and the Andaman Islands), Thailand, Malayisa (Peninsular and Sabah, Borneo, the Philippine island of Palawan, and the Indonesian islands of Timor and Flores (100% support). The short sequences from Mizoram, India and Myanmar (Table 1) most closely matched those of the *A. dorsata dorsata* group and were clearly distinct from the A. d. South India group.

Unfortunately, although this analysis shows four well-supported lineages within *A. dorsata* in the broad sense, it does not resolve branching patterns among the four lineages. Although color, morphometric and morphological differences among *A. d. dorsata, A. d. binghami* and *A. d. breviligula* have been found (e.g., Maa, 1953), there are no obvious morphological differences between A. dorsata South India and the widespread *A. d. dorsata*. In their study of *A. laboriosa* and *A. dorsata* in India, Kitnya et al. (2022) examined specimens of *A. dorsata* from Arunachal Pradesh in the extreme northeast of India and Karnataka (specifically Bangalore) in south India. In a dendogram displaying morphometric similarity of the samples, the south India specimens did not form a discrete cluster but were mixed in among the specimens from northeast India. However, in their phylogenetic analysis (using a 500 bp fragment of the mitochondrial COI gene), the samples from Arunachal Pradesh and south India formed two separate clades with 99% and 100% bootstrap support, respectively. One earlier publication (Sharma and Thakur, 1999) reported a morphometric differentiation between the giant bees of the Doon valley and other populations of *A. dorsata dorsata* in India; among other characteristics, it was generally larger than other populations. However, this is very unlikely to represent A. d. South India as the Doon Valley is in Uttar Pradesh, at the extreme north of India.

To our knowledge, there is no information on the geographical distribution of *A. d. dorsata* and A.d. South India. Although giant honeybees have been collected at points between southern and northern India, at the moment the only way to tell the two apart is by genetic testing.

## Discussion

In this study we show that the giant honeybee *Apis dorsata* in the broad sense includes four genetically distinguishable lineages: *A. dorsata dorsata, A. d. binghami, A. d. breviligula* and A. dorsata South India. Regardless of whether the four lineages within *A. dorsata* in the broad sense merit species status, recognition and continued investigation of these groups are important, particularly for study of honeybee biogeography and migratory behavior of giant bees, and for conservation programs.

The distinctive nature of the giant honeybees of the oceanic Philippine islands (i.e., those islands that were not part of the Sundaland region) and of Sulawesi and adjacent small islands has long been recognized (e.g., Maa, 1953) The fact that isolated island populations show traits distinct from those of mainland populations and from each other is perhaps not surprising.

What is more surprising is the presence of a genetically distinct giant honeybee population in southern India, along with the more widespread *A. dorsata dorsata* in northern India. However, a broader view of Indian *Apis* shows that this pattern has appeared there more than once. India is also home to two cavity-nesting bees, the yellow or plains bee, and the hill or black bee (Ruttner, 1988 and references cited therein; Bhatta et al., 2020). According to Engel (2002) the yellow or plains bee corresponds to *A. cerana indica* Fabricius while the black or hill bee corresponds to *A. cerana cerana* Fabricius. Genetic evidence collected over the past three decades (e.g., mitochondrial COI, COII and non-coding sequences, Smith, 1991, Smith and Hagen, 1996; mitochondrial and nuclear gene sequences, Lo et al., 2010; and genomic SNPs, Su et al., 2023) support species status of the yellow Indian bee, as proposed by Lo et al., 2010 and Su et al., 2023. The dwarf honeybee, *Apis florea*, consists of two distinct groups revealed by mitochondrial COI-COII sequences and nuclear SNPS (Smith, 2011, Su et al., 2023, and Figure 1 of this study). These are an eastern lineage including populations from Thailand eastwards, and a western lineage including populations from India westward. The “switchover” is apparently in the poorly sampled region from northeastern India through Bangladesh and Myanmar.

Why does India have a distinct variety of cavity-nesting bee, *A. cerana indica* along with *A. cerana cerana*, and a distinct south Indian variety of giant honeybee along with *A. dorsata dorsata* in northern India? And why does it have a different *A. florea* from that in eastern Asia? Answering these questions requires (1) information on the ranges of the species and putative species of *Apis* in India, particularly the distributions of the yellow and black cavity-nesting bees, and A. d. South India and *A. dorsata dorsata*, and (2) a phylogeny that resolves the branching patterns of the four *A. dorsata* lineages. A robust phylogeny would provide information on the order and timing of diversification events. A time calibrated phylogeny could suggest specific geological and climatic events that could have promoted diversification, and help us determine if the dwarf, giant, and cavity nesting lineages responded to historical events with similar patterns of diversification.

Recognition of these four putative or redefined species is important for pollinator conservation planning. Each exhibits migratory behavior, typically in predictable annual responses to seasonal patterns of rainfall and resource availability (e.g., Dyer et al., 1994) or in response to erratically occurring masting events in which forest trees produce a superabundance of blossoms and resources (Itioka et al., 2001). Migrating bees appear to show fidelity to their nesting sites at either end of the migratory route (Neumann et al., 2000, Paar et al., 2000). This alone means that protecting a giant bee’s nesting and foraging habitat means protecting more than one location. Migrating colonies of giant bees may travel distances that require “rest stops” to forage Recent work by Robinson (2012, 2021) has shown that migrating *A. d. dorsata* in Thailand also make use of “traditional” rest stops, where they forage for food and water for variable lengths of time before continuing their journey. These rest stops are also likely to be crucial for successful migration. The vast majority of giant bee migration research has been carried out on populations that our study would place in *A. d. dorsata* (for example, Dyer & Seeley, 1994, Kahono et al., 1999, Neumann et al., 2000, Paar et al., 2000, Itioka et al., 2001, Rattanawannee et al., 2013), which is not surprising, as it is the most widespread. Koeniger & Koeniger (1980) investigated giant bee migration in Sri Lanka; though we have not sampled any giant bees from Sri Lanka, we predict that they are part of the A. d. South India clade, based on the fact that the south Indian plains bee, *A. c. indica*, is also found in Sri Lanka. At least one set of observations has been made on the migration of *A. d. binghami* in Sulawesi (Nagir et al., 2016). Morse & Laigo (1969, cited in Robinson, 2021) reported that *A. d. breviligula* in the Philippines does not migrate.

Our results suggest avenues for additional research, particularly regarding Indian populations. What is the range of the South Indian giant bee, and what are its migration patterns? Are the ranges of the southern Indian cavity-nesting “plains bee” (currently *A. cerana indica*) congruent with those of the southern Indian giant bee, implying similar biogeographic history? Do they differ in morphology, behavior or ecology from *A. d. dorsata*? And of course, will behavioral and genetic study of giant honeybees from a greater portion of their ranges (e.g., Cao et al., 2012b) reveal more diversity?

## Acknowledgements

We would like to thank the many people who have helped us in the field and by collecting and donating specimens:

Ahmed Al-Ghamdi, Nicola Bradbear, Fred Dyer, Steven Goodman, the late Randall Hepburn, Ben Oldroyd, Jurgen Paar, the late Herman Pechhacker, Stephen Petersen, the late Stefan Reyes, Benny Shalmon, Yong-Chao Su, and especially Gard Otis, who helped many bee researchers begin their studies of Asian honeybees.

A very large portion of this work was completed by Sarah Cluff in partial fulfilment of the requirements for an Honors thesis and Bachelor of Science (Honors) degree in Biological Sciences at the University of Kansas.

## Literature cited

Arias, M. C., and Sheppard, W. S. (2005). Phylogenetic relationships of honeybees (Hymenoptera: Apinae: Apini) inferred from nuclear and mitochondrial DNA sequence data. Molecular Phylogenetics and Evolution 37: 25–35. https://www.sciencedirect.com/science/article/abs/pii/S1055790305000667

Bhatta, C. P., Reddy, M. S., and Smith, D. R. (2020). Scientific note: Varroa jacobsoni and V. destructor on hill and plains strains of Apis cerana in southern India. Apidologie, 51:391–394. https://link.springer.com/article/10.1007/s13592-019-00723-7

Cao L-F., Zheng H-Q., Chen X., Niu D-F., Hu F-L., & Hepburn, H. R. (2012a). Multivariate morphometric analyses of the giant honeybees, Apis dorsata F. and Apis laboriosa F. in China. Journal of Apicultural Research 51: 245–251. https://www.tandfonline.com/doi/abs/10.3896/IBRA.1.51.3.05

Cao, L-F., Zheng, H-Q., Hu, C-Y., He, S-Y., Kuang, H-O. and Hu, F-L. (2012b). Phylogeography of the giant honeybee Apis dorsata (Hymenoptera: Apidae) from China and neighbouring Asian areas. Annals of the Entomological Society of America 2: 298–304. 10.1603/AN11104

Cornuet, J-M., Garnery, L., and Solignac, M. (1991). Putative origin and function of the intergenic region between COI and COII of Apis mellifera L. Mitochondrial DNA. Genetics 128: 393–403. https://academic.oup.com/genetics/article/128/2/393/6006820

Crozier, R. H., and Crozier, Y. C. (1993). The mitochondrial genome of the honeybee Apis mellifera: complete sequence and genome organization. Genetics 133: 97–117. https://academic.oup.com/genetics/article/133/1/97/6009209

Dyer, F. C. and Seeley, T. H. (1994). Colony Migration in the tropical honeybee Apis dorsata F. (Hymenoptera: Apidae). Insectes Sociaux 41:129–140. https://link.springer.com/article/10.1007/BF01240473

Engel, M. S. (1999). The taxonomy of recent and fossil honeybees (Hymenoptera: Apidae; Apis). Journal of Hymenoptera Research 8: 165–196.

Engel, M. S. (2002). The honeybees of India, Hymenoptera: Apidae. Journal of the Bombay Natural History Society 99(1): 3–7.

Hall, H. G., & Smith, D. R. (1991). Distinguishing African and European honeybee matrilines using amplified mitochondrial DNA. Proceedings of the National Academy of Science USA 88:4548–4552. https://www.pnas.org/doi/abs/10.1073/pnas.88.10.4548

Hepburn, H.R., Smith, D.R., Radloff, S. E., and Otis, G.W. (2001). Infraspecific categories of Apis cerana: morphometric, allozymal and mtDNA diversity. Apidologie 32:3–23. DOI: 10.1051/apido:2001108

Itioka, T., Inoue, T., Kaliang, H., Kato, M., Nagamitsu, T., Momose, K., Sakai, S., Yumoto, T., Mohamad, S. U., Hamid, A. A., and Yamane S. (2001). Six-Year Population Fluctuation of the Giant Honeybee Apis dorsata (Hymenoptera: Apidae) in a Tropical Lowland Dipterocarp Forest in Sarawak. Annals of the Entomological society of America 94: 545–549. 10.1603/0013-8746(2001)094[0545:SYPFOT]2.0.CO;2

Kitnya, N., Prabhudev, M. V., Bhatta, C. P., Pham, T. H., Nidup, T., Megu, K., Chakravorty, J., Brockmann, A., and Otis, G. W. (2020). Geographical distribution of the giant honeybee Apis laboriosa Smith, 1871 (Hymenoptera, Apidae). Zookeys 951: 67–81. Doi: 10.3897/zookeys.951.49855

Kitnya, N., Otis, G.W., Chakravorty, J., Smith, D.R., and Brockmann, A. (2022). Apis laboriosa confirmed by morphometric and genetic analyses of giant honeybees (Hymenoptera, Apidae) from sites of sympatry in Arunachal Pradesh, Northeast India. Apidologie 53: 47, 17 pp. 10.1007/s13592-022-00956-z

Koeniger, N., and Koeniger, G. (1980). Observations and experiments on migration and dance communication of Apis dorsata in Sri-Lanka. Journal of Apicultural Research 19: 21–34. 10.1080/00218839.1980.11099994

Kohono, S., Nakamura, K., and Amir, M. 1999. Seasonal migration and colony behavior of the tropical honeybee Apis dorsata F. (Hymenopter: Apidae). Truebia 31: 283–297. DOI: 10.14203/treubia.v31i3.611

Kumar, S., Stecher, G., and Tamura, K. (2016). MEGA7: Molecular Evolutionary Genetics Analysis version 7.0 for bigger datasets. Molecular Biology and Evolution 33: 1870–1874. 10.1093/molbev/msw054

Lo, N., Gloag, R. S., Anderson, D. L., and Oldroyd, B. P. (2010). A molecular phylogeny of the genus Apis suggests that the giant honeybee of the Philippines, A. breviligula Maa, and the plains honeybee of southern India, A. indica Fabricius, are valid species. Systematic Entomology 35: 226–233. 10.1111/j.1365-3113.2009.00504.x

Maa, T. C. (1953). An inquiry into the systematics of the tribus Apidini or honeybees (Hymenoptera). Treubia 21: 525–640.

McEvoy, V. M., and Underwood, B. A. (1988). Drone and species status of the Himalayan honeybee, Apis laboriosa (Hymenoptera: Apidae). Journal of the Kansas Entomological Society 61: 246–249. http://www.jstor.org/stable/25084995

Morse, R. A., and Laigo, F. M. (1969). Apis dorsata in the Philippines. Philippine Association of Entomologists, Laguna, Philippines.

Nagir, M. T., Atmowidi, T., and Kahono, S. (2016). The distribution and nest–site preference of Apis dorsata binghami at Maros Forest, South Sulawesi, Indonesia. Journal of Insect Biodiversity 4(23): 1–14. DOI: 10.12976/jib/2016.4.23

Neumann, P., Koeniger, N., Koeniger, G., Tingek, S., Kryger, P., and Moritz, R. F. A. (2000). Home–site fidelity in migratory honeybees. Nature 406: 474–475.

Paar, J., Oldroyd, B. P., and Kastberger, G. (2000). Giant honeybees return to their nest sites. Nature 406: 475.

Raffiudin, R., and Crozier, R. H. (2007). Phylogenetic analysis of honeybee behavioral evolution. Molecular Phylogenetics & Evolution 43: 543–552. 10.1016/j.ympev.2006.10.013

Rattanawannee, A., Chanchao, C., Lim, J., Wongsiri, S., and Oldroyd, B. P. (2013). Genetic structure of a giant honeybee (Apis dorsata) population in northern Thailand: implications for conservation. Insect Conservation and Diversity 6:38–44. 10.1111/j.1752-4598.2012.00193.x

Robinson, W. (2012). Migrating giant honeybees (Apis dorsata) congregate annually at stopover site in Thailand. PLoS ONE 7(9): e44976. doi:10.1371/journal.pone.0044976

Robinson, W. (2021). Surfing the Sweet Wave: migrating giant honeybees (Hymenoptera: Apidae: Apis dorsata) display spatial and temporal fidelity to annual stopover site in Thailand. Journal of Insect Science 21(6):1–12. 10.1093/jisesa/ieab037

Ruttner, F. (1988). Biogeography and taxonomy of honeybees. Springer, Berlin, Germany.

Sakagami, S. F., Matsumura, T. and Ito, K. (1980). Apis laboriosa in Himalaya, the little-known world largest honeybee (Hymenoptera, Apidae). Insecta Matsumurana 19: 47–77. http://hdl.handle.net/2115/9801

Sharma, V. and Thakur, M. L. (1999). Morphometric characterization of giant honeybee Apis dorsata Fab. (Hymenoptera: Apidae) from Doon Valley. Ann. For. 7(1): 125–135.

Smith, D. R. (1991). Mitochondrial DNA and honeybee biogeography. pp. 131-176 in: Diversity in the Genus Apis, DR Smith (ed.), Westview Press, Boulder, CO.

Smith, D. R. (2011). Asian honeybees and mitochondrial DNA. pp. 69-93 in: Honeybees of Asia, HR Hepburn & SE Radloff (eds.) Springer Verlag, New York. 681 pp.

Smith, D. R., and Hagen, R. H. (1996). The biogeography of Apis cerana as revealed by mitochondrial DNA sequence data. Journal of the Kansas Entomological Society 69(4) suppl., 294–310. https://www.jstor.org/stable/25085726

Su, Y-C., Chiu, Y-F., Warrit, N., Otis, G. W. and Smith, D. R. (2023). Phylogeography and species delimitation of the Asian cavity-nesting honeybees. Insect Systematics and Diversity 7(4), 5; 1–10 10.1093/isd/ixad015

